# Adaptation of the eyes of grass puffer (*Takifugu niphobles*) to the riverine and marine environments

**DOI:** 10.1101/2022.04.28.489969

**Authors:** Misa Osada, Yohey Terai

## Abstract

Many marine fish species migrate to rivers, but little is known about whether these species switch their vision when inhabiting rivers or adapt their vision to the environment of both rivers and the sea. Grass puffer (*Takifugu niphobles*) is a marine fish species frequently migrating to rivers. In this study, we investigated grass puffers from riverine and marine populations and analyzed the gene expression in their eyes. The phylogeny and population genetics of riverine and marine grass puffers indicated that riverine and marine grass puffers are from the same population. Gene expression levels by high-throughput RNA sequencing indicated no differences in the expression patterns of vision-related genes in marine and riverine grass puffers. This result indicates that the adaption of their visual system to both marine and riverine environments rather than switching the expression of vision-related genes. Additionally, riverine grass puffers increase the expression levels of heat shock proteins and related genes. These genes showed higher expression in riverine grass puffer than in other marine and river pufferfish species, suggesting that the grass puffer individuals adapt to the environmental difference when they migrate to the river by increasing the expression levels of heat shock protein and related genes.

## Introduction

The environments in the rivers and the sea differ in many aspects, including nutrient, salinity, light intensity, and wavelength of light. For example, in freshwater with low transparency, water with micro-particles scatters short-wavelength light, and the light in turbid water is dominated by long wavelengths ^1-3^. In contrast, long-wavelength light is absorbed by water, and short-wavelength light is predominant in the clear ocean ^4^. Since fish species inhabit this heterogeneous light environment, they have adapted their vision to the different light environments ^2,3,5,6^. Accordingly, fish vision is known as a prime example of adaptive evolution.

Vertebrate visual pigments consist of light-absorbing chromophores and the protein component, opsin ^7^. Spectral sensitivity is determined by the type of chromophore [with 11-*cis* retinal (A1-) or 11-*cis* 3-dehydroretinal (A2-derived retinal)] and by interactions between chromophores and amino acid residues of opsin ^8^. The replacement of A1- with A2-derived retinal in the visual pigments shifts the absorption to a longer wavelength ^9,10^.

The enzyme CYP27c1 is known to be involved in the switch from A1 to A2, and the expression level of this gene correlates with the amount of A2 ^11,12^. Fish species use A1 and A2 retinal, with freshwater fish mainly using A2 and marine fish using A1 retinal ^13^. A1 and A2 retinal are known to be differentially used at different life cycle stages of the life cycle. Salmon use A1 retinal when living in the sea, and switch to A2 when they migrate up the river to spawn ^14^. In lamprey, A1 and A2 are differentially used for juveniles and upstream migrating adult lamprey ^11^. Marine fish species migrating to the river use both A1 and A2 retinal ^13^. This usage is a characteristic of both riverine and marine environments. However, little is known about whether these species mainly use A2 retinal when they live in rivers or whether they have both A1 and A2 retinal in both rivers and the sea. Furthermore, little is known about whether these species express different levels of opsin genes in the rivers and the sea.

Grass puffer (*Takifugu niphobles*) is a marine fish species frequently migrating to rivers. Grass puffers are abundant in rivers and the sea during the same season. Fish with ancestry in marine fish and colonized freshwater have been reported to have adapted to freshwater by duplicating specific genes ^15,16^. However, the grass pufferfish would not be a colonizing fish because they cannot live in freshwater for long periods ^17^. Grass puffers possess both A1 and A2 retinal ^13^, therefore they seem to migrate to rivers for short periods. Hence, it is unclear how they adapt to different river environments compared to the sea and whether the populations in the rivers and the sea are the same. In this study, we analyzed the phylogenetic and population genetics of riverine and marine populations of grass puffer. Also, we analyzed gene expression levels in the eye of grass puffer from both riverine and marine populations. In addition, the gene expression levels in the eyes of marine and riverine pufferfish species were compared with those of riverine and marine grass puffers to show how grass puffers inhabit the river and marine environments.

## Result

### No genetic differentiation between the riverine and marine populations of *T. niphobles*

We collected riverine grass puffers 1.3 km upstream from the mouth of Tagoe River, Kanagawa, Japan (Fig. S1, Table S1). Based on our observations, the grass puffers are not seen in the rivers during midwinter (mid-January through the end of February). In contrast, we visually observed many grass puffer individuals in the rivers from March through December, even at low tide. We collected Marine grass puffers on a beach 10 km from the mouth of Tagoe River. Since this beach is more than 3 km from the nearest river, we expected less direct migration of the river grass puffers. We measured salinity and light environments at the Tagoe River and Tateishi Park beach to verify the differences between the river and the sea environments (Fig. S2).

As a first step, we investigated the genetic relationships between the river and marine populations of the grass puffer. Single nucleotide polymorphisms (SNPs) were extracted from short reads of grass puffers from the riverine and marine populations (see Methods) mapped to coding regions of a reference genome (*Takifugu rubripes*). We calculated Fst values using a total of 172,214 SNPs. The mean Fst value between riverine and marine populations was almost 0 (-0.00026635), indicating no genetic differentiation between these two populations.

*T. rubripes* and *T. poecilonotus* are marine species in the same genus as grass puffer. SNPs were extracted from individuals of these species collected from the sea (Fig. S1, Table S1) using the same method as grass puffer individuals (see Methods). The SNPs were combined with the grass puffer SNPs, and a phylogenetic tree was constructed by the ML method using *C. rivulata* and *T. nigroviridis* as out-group species with a total of 116,941sites. Phylogenetic trees showed that grass puffer formed a monophyletic group with no population differentiation (Fig. 1). We then performed a population genetic analysis with 275,155 sites. Using the same data set, a principal component analysis (PCA) showed similar results, with no differentiation between the riverine and marine populations of grass puffer (Fig. 2A). Furthermore, ADMIXTURE analysis using the same data set showed that the grass puffers formed either a single cluster (K=2, 3) or multiple clusters (K=4-6), but the riverine and marine populations did not differentiate (Fig. 2B). These results indicate that riverine and marine grass puffers are from the same population. Grass puffer is known to spawn simultaneously in the surf on the beach in June, and subpopulations are presumed to mix at this time, which may prevent genetic differentiation.

**Fig. 1.**
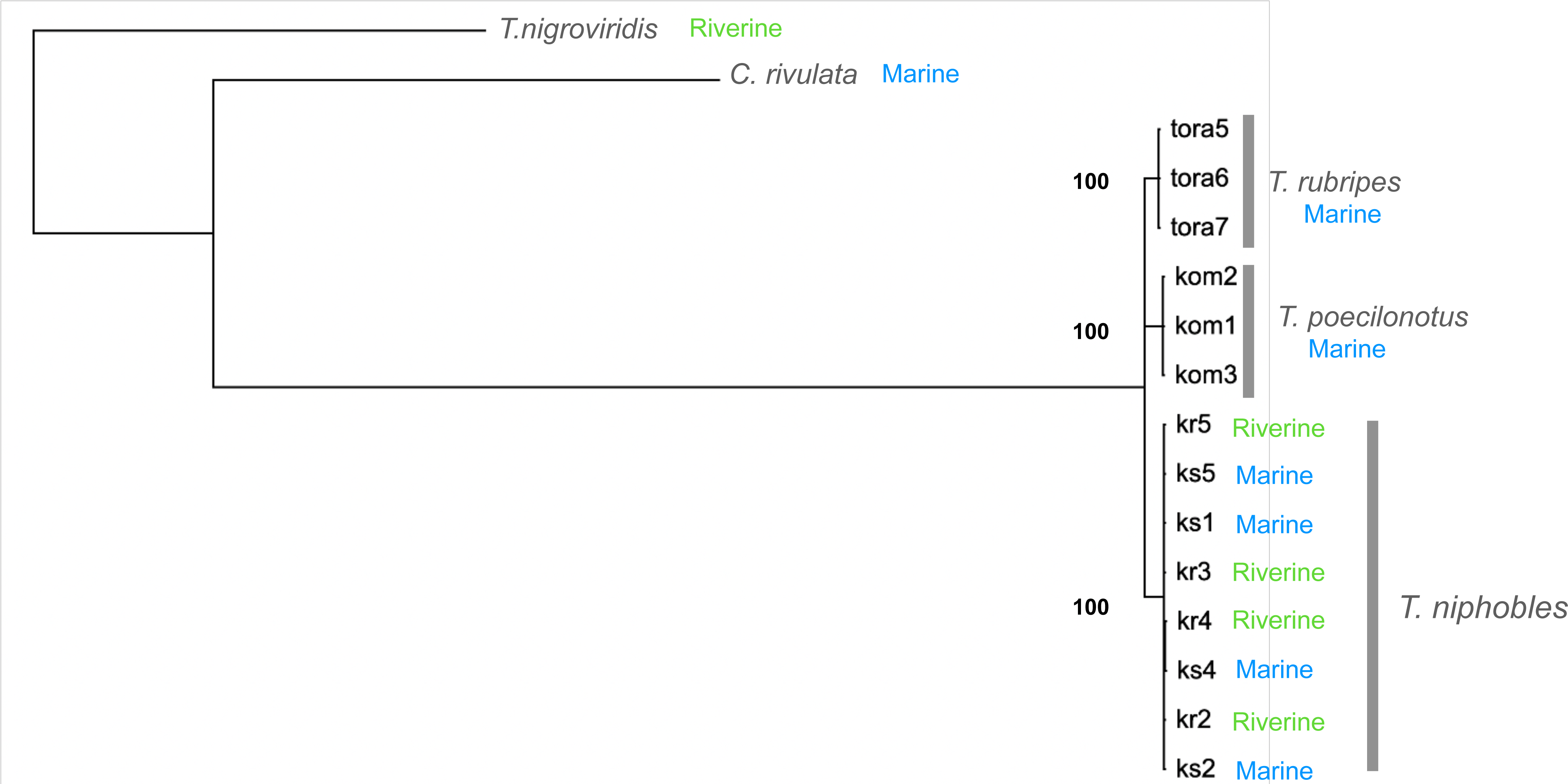
Phylogenetic relationships between *Takifugu* species. Maximum likelihood tree based on 266,954 SNPs. Node labels indicate bootstrap replicates.

**Fig. 2.**
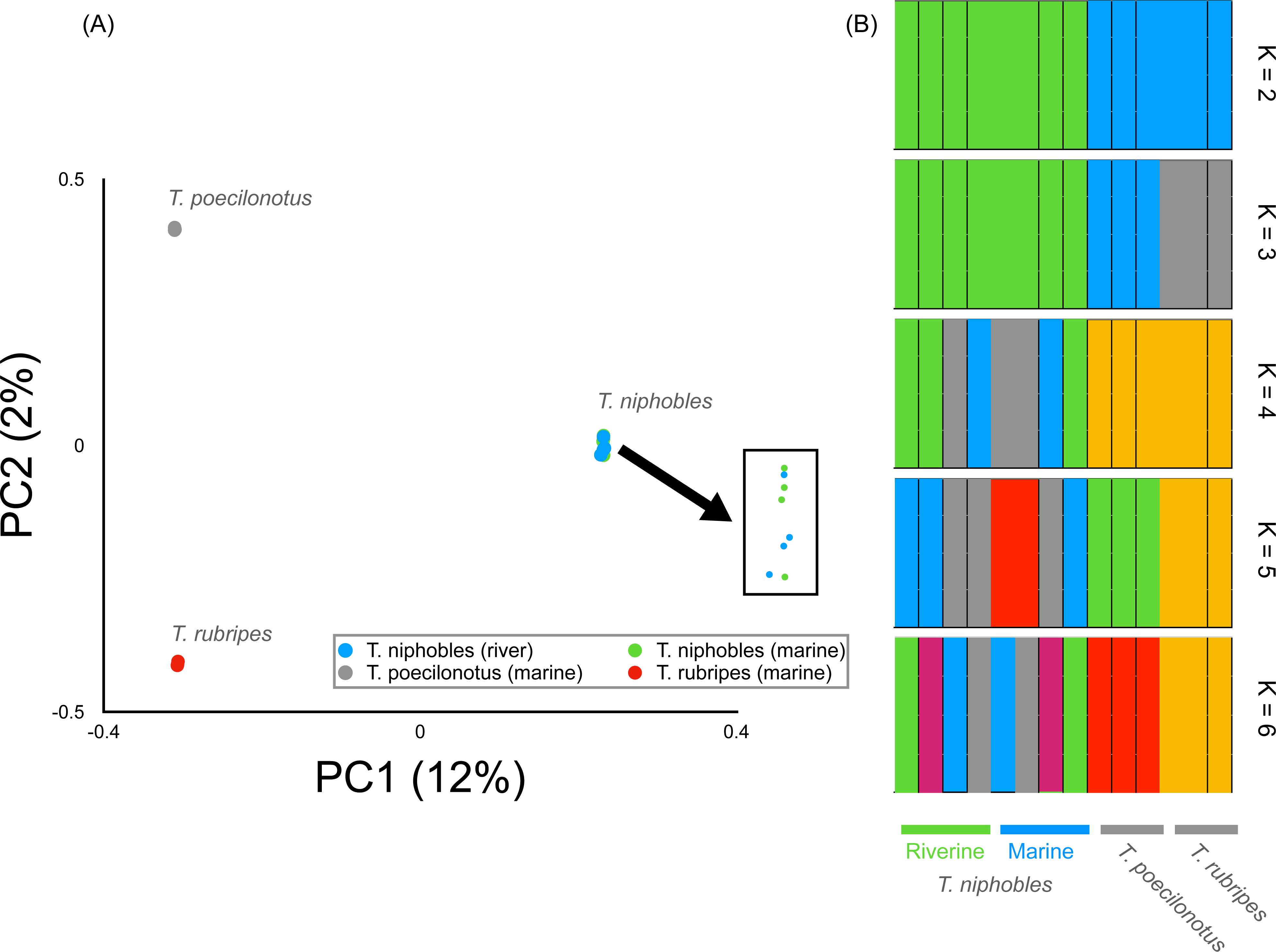
Genetic relationships between *Takifugu* species. Principal Components Analysis (PC1 versus PC2) based on 266,954 SNPs (A). Colored circles correspond to the names of species or populationss in the panel. (B) ADMIXTURE results based on the same SNP data with (A) for K = 2-6.

### Expression differences in the eyes between the riverine and marine populations of *T. niphobles*

We performed RNA sequencing to compare gene expression levels in the eyes of grass puffers from the riverine and marine populations. Total RNAs were extracted from eyes, and 10-15 Gb sequences were determined from four individuals, each of grass puffers from the riverine and marine populations. These high-throughput sequences were mapped to a reference genome sequence of *T. rubripes* to compare gene expression levels between the riverine and marine populations of grass puffers. As a result, we found 25 differentially expressed genes with statistical significance (Fig. 3, Table S2). Among the 25 genes, six genes were highly expressed in the riverine population, and 19 genes were highly expressed in the marine population (Fig. 3, Table S2). In this study, we focused on six of the 25 genes highly expressed in the riverine population to reveal how marine grass puffer individuals can inhabit a freshwater environment. Three of these were heat shock protein genes (Hsp 9, 47, 90) ^18,19^. unc45b has also been reported to be part of a chaperone system that interacts with HSP90 ^20^.

**Fig. 3.**
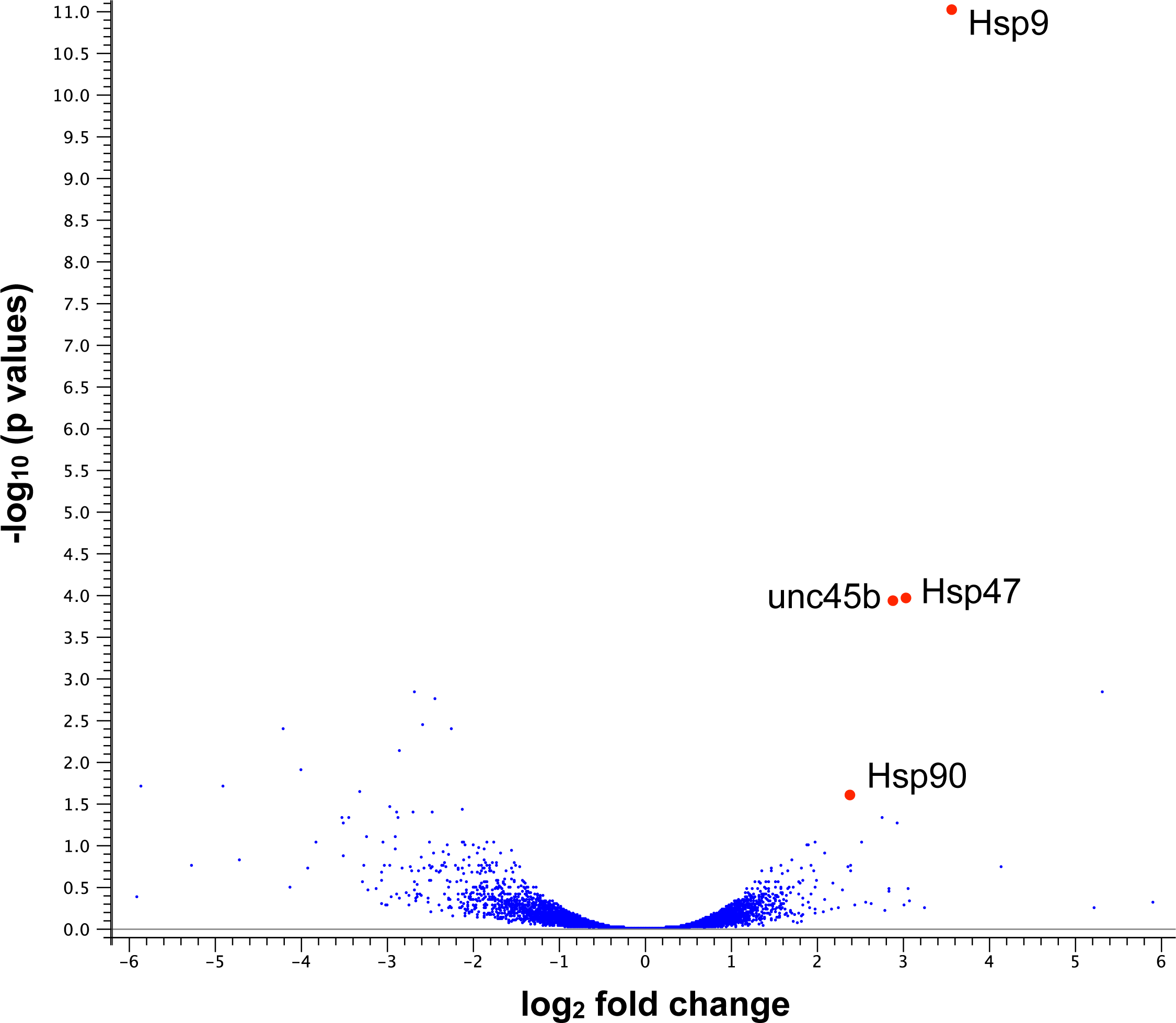
A volcano plot for the gene expression differences between the riverine and marine grass puffer populations. Each blue dot represents the gene expressed in the eye. The log2 fold changes of average TPM values of the marine grass puffer population compared to those of the riverine grass puffer population and the -log10 p-values resulting from a t-test comparing average TPM values between the riverine and marine grass puffer populations for each gene are plotted on the x- and y-axis, respectively.

Next, we examined whether the expression levels of genes related to vision varied between the riverine and marine populations. The photosensitivity of visual pigments is tuned by the amino acid sequence of opsins and the type of chromophores ^8^. Therefore, we first searched for the opsin genes in the genome of *T. rubripes* and CYP27c1 gene for switching chromophores. The color opsins, blue-sensitive (*SWS2)*, green-sensitive (*RH2)*, and long-wavelength sensitive (*LWS)* opsin genes were annotated in the genome of *T. rubripes*. In addition, the scotopic opsin, *RH1* gene, and *CYP27c1* were present in the genome. Comparisons of gene expression levels of the four opsin genes and *CYP27c1* between the river and marine populations of grass puffer showed no difference with statistical significance (Fig. 4). In addition, the amino acid sequences of four opsins were identical between the two populations.

**Fig. 4.**
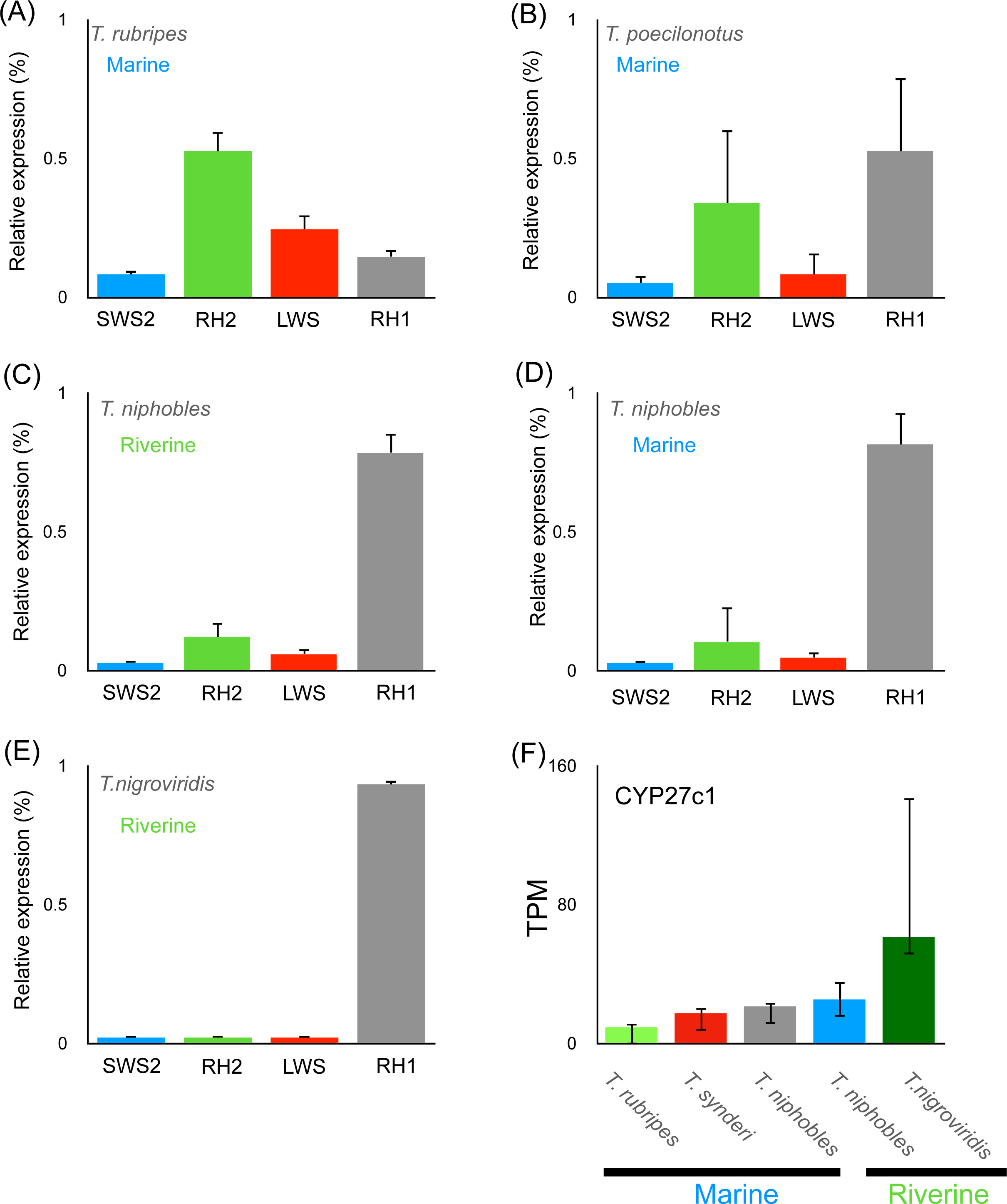
Expression of opsin and CYP27c1 genes. The relative expression of opsin genes in (A) *T. rubripes*, (B) *T. poecilonotus*, (C) riverine population of grass puffer, (D) marine population of grass puffer, and (E) *T. nigroviridis*. (F) the expression of CYP27c1 gene.

### Comparisons of gene expression between species in Tetraodontidae

The opsin repertoire was examined for *T. rubripes* and *T. poecilonotus*, as well as *Tetraodon nigroviridis*, which is a Tetraodontidae species inhabiting rivers and ponds in Southeast Asia ^21^. We also searched for the *SWS2*, *RH2*, *LWS*, and *RH1* genes in the assembled RNA sequences of *T. rubripes*, *T. poecilonotus,* and *T. nigroviridis*, and found the homologs of these genes. Therefore, we infer that the genus *Takifugu*, including grass puffer, and *T. nigroviridis* have an opsin repertoire of *SWS2*, *RH2*, *LWS*, and *RH1* and use a trichromatic color vision.

We focused on four opsin genes, CYP27c1, and genes with higher expression levels in the riverine population than in the marine population of the grass puffer and compared the expression levels with those of *T. rubripes*, *T. poecilonotus*, and *T. nigroviridis*. The relative expression of *RH*2 in marine *T. rubripes* and *T. poecilonotus* exceeded 30% of total opsin expressions, and the expression of *SWS2* and *LWS* was also relatively higher than that of the grass puffer (Fig. 4A, 4B). Therefore, these two marine species primarily express color opsin genes. In contrast, the relative expression of *RH1* in riverine *T. nigroviridis* exceeded 95% of total opsin expressions, indicating that a scotopic opsin gene was predominantly expressed in this species (Fig. 4E). In the grass puffer, *RH1* was predominantly expressed, but less than in *T. nigroviridis*, with 67% and 84% in the riverine and marine populations, respectively (Figs. 4C-4E). *SWS2*, *RH2,* and *LWS* were highly expressed in grass puffer than in *T. nigroviridis*. *CYP27c1* was expressed at lower levels in the marine species *T. rubripes* and *T. poecilonotus* than in the grass puffer and higher levels in *T. nigroviridis* compared to the grass puffer (Fig. 4F). Among the six genes with higher expression levels in the riverine population than in the marine population of the grass puffer, we focused on four highly expressed genes with (TPM > 5). These four genes expressed higher levels in the river population of the grass puffer than in the other species (Fig. 5).

**Fig. 5.**
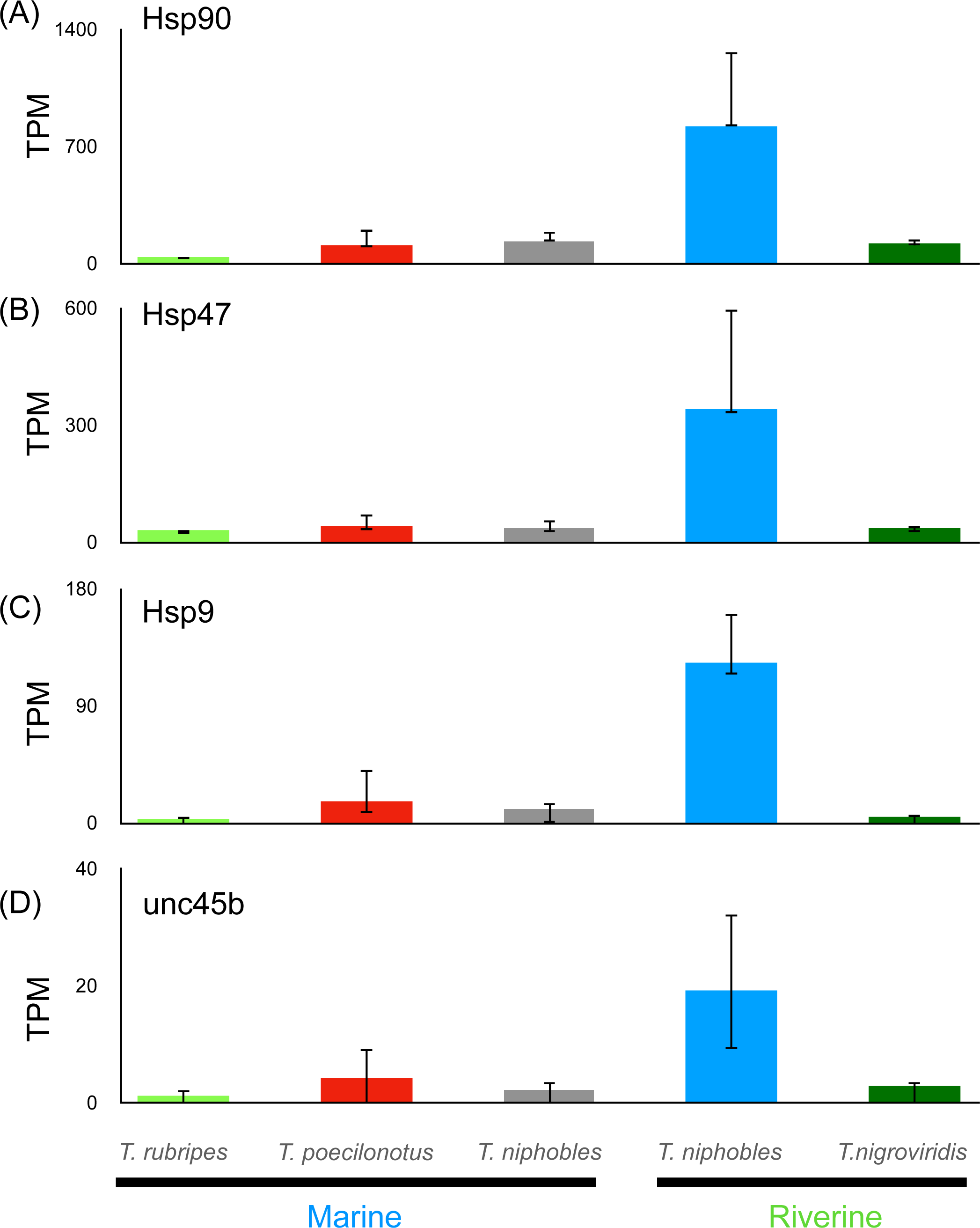
Expression of heat shock protein and related genes. The expression of (A) Hsp 90, (B) Hsp47, (C) Hsp 9, and (D) unc45b in *T. rubripes*, *T. poecilonotus*, riverine and marine populations of grass puffer, and *T. nigroviridis*. Asterisks indicate statistical significance (p < 0.05 after Bonferroni correction) of differences in expression levels between riverine grass puffer and *T. rubripes* and *T. poecilonotus*.

## Discussion

When marine fish migrate to a river, they face a riverine environment that differs from the sea. Therefore, we expected that marine fish that migrate to the river might respond to a different environment, and this study investigated mRNA expression in the eyes of the grass puffer, which inhabits both the river and the sea. As for vision, the repertoire of opsin genes and their expression levels were comparable between the riverine and marine populations of the grass puffer. The expression levels of *CYP27c1*, representing the amount of A2 retinal, were also comparable. Thus, grass puffers are presumed to have the same vision usage in both marine and riverine environments. In contrast, *RH1* was predominantly expressed in the river species *T. nigroviridis*, while cone opsins were predominantly expressed in the marine species *T. rubripes* and *T. poecilonotus*. In the expression of the opsin genes of all species used in this study, the grass puffer showed an expression pattern intermediate between that of marine and river species. The expression of *CYP27c1* was also intermediate between marine and riverine species. According to these results, we inferred that the usage of the visual system in the grass puffers is the same in both the river and the sea. In other words, the grass puffers adapt their visual system to two environments, the river and the sea, rather than switching their visual system by changing the expression of vision-related genes depending on the ambient environments.

Unlike the vision-related genes, the expression levels of the six genes were significantly upregulated in the riverine grass puffer population compared to the marine population. In the four highly expressed genes among these six genes, the expression levels were upregulated in the riverine grass puffer population compared to the marine *T. rubripes* and *T. poecilonotus*. Interestingly, the expression levels of these four genes were higher in the riverine grass puffer population than in the riverine *T. nigroviridis*. These four genes were heat shock proteins (Hsp), Hsp 90, Hsp47, Hsp9, and unc45b. Hsp functions in protein folding, maintenance of structural integrity, and proper regulation for a wide range of proteins ^18^. Hsp90 has been reported to play a role in lens formation in the eye ^22^. In addition, unc45b is known to interact with Hsp90 and is involved in lens formation ^20^. Hsp47 is a molecular chaperone specific to collagen and plays an essential role during ocular development, especially in corneal morphogenesis ^23^. Considering that these three chaperone-related genes function in lens and cornea formation, these genes may be involved in normal eye formation and maintenance in freshwater environments with lower salt concentrations than seawater in grass puffer. *T. nigroviridis* also inhabits the river, but the expression levels of these three genes were not upregulated. This species adapts to the river, thus the riverine environment is probably not a major stressor for them. Hsp9 has been proposed to be involved in the immune response against pathogenic bacteria ^19^. The grass puffer individuals that have migrated to the river likely face a different type of pathogenic bacteria than those in the sea and have upregulated expression of Hsp9.

Taken together, we hypothesize that the grass puffer individuals adapt to the environmental difference (low salinity and pathogenic bacteria) that they face when they migrate to the river by increasing the expression levels of heat shock protein and related genes.

### Conclusion

In this study, we investigated how the grass puffer inhabits two different environments, the river and the sea, based on gene expression in the eyes. Opsin and *CYP27c1* genes were expected to be adapted to riverine and marine environments by making their expression patterns intermediate between riverine and marine species. On the other hand, the high expression of *Hsp* genes in the riverine grass puffer population suggests that they adapt to the river environment by expressing heat shock proteins. Future studies of other marine fish migrating to rivers will reveal whether other fish species share these two adaptive strategies to different environments.

## Methods

### Measurement of salinity and light environments

We measured salinity at the Tagoe River and Tateishi Park (Fig. S1). In the Tagoe River, we collected water from a depth of 50 cm every hour from low to high tide at the point where we collected the grass puffers and measured salinity using a hydrometer (SpectrumBrands Japan, Kanagawa Japan) (Fig. S2A). In Tateishi Park, we measured salinity only once since salinity does not change with tides. We measured the wavelength of light in the horizontal direction at a depth of 50 cm with an Ocean Optics spectrophotometer (Ocean Photonics, Tokyo Japan) at the sites where we collected the grass puffers in the Tagoe River and Tateishi Park (Figs. S1 and S2 B).

### Sampling

Grass puffers were collected from Tagoe river and Tateishi park (Fig. S1) by fishing. *T. rubripes* and *T. poecilonotus* individuals were collected from Tokyo bay (Fig. S1) by fishing*. C. rivulata* was collected from Tateishi park (Fig. S1) by fishing. *T. nigroviridis* individuals were obtained from tropical fish dealer. The information of each species used in this study are listed in Table S1. The guidelines for experimental animal management of SOKENDAI were followed throughout the study. The Institutional Animal Care and Use Committee of SOKENDAI approved the animal protocols and procedures.

### RNA sequencing and comparison of expression levels

Total RNAs were extracted from eyes of *C. rivulata, T. rubripes*, *T. poecilonotus*, *T. nigroviridis*, and grass puffer using TRIzol RNA Isolation Reagent (Thermo Fisher Scientific, Waltham, MA, USA) according to the manufacturer’s instructions. RNA libraries were constructed using the NEBNext Poly(A) mRNA Magnetic Isolation Module and the NEBNext Ultra RNA Library Prep Kit for Illumina (New England Bio Labs, Ipswich, MA) following the manufacturer’s instructions. Short cDNA sequences (paired-end 150 bp) were determined from the libraries using the Illumina HiseqX platform (RNA-seq). RNA-seq reads from each of three or four individuals of *T. rubripes*, *T. poecilonotus*, and grass puffer (10-15 Gbp) were trimmed to remove adaptor sequences and mapped to the reference genome of *T. rubripes*. Reads showing similarity (90%) with 90% read lengths were mapped to the reference genome and expression levels were calculated using CLC Genomics Workbench ver. 21 (https://www.qiagenbioinformatics.com/). TPM (Transcript Per Million) was used for normalized expression values. The average TPM of the four individuals of the grass puffer were compared. Differential expression was assessed using the TMM (trimmed mean of M values) method ^24^ implemented in the CLC genomics workbench using default parameters. False Discovery Rate (FDR) correction was applied to the calculated p-values and differentially expressed genes with corrected p-value < 0.05 were considered significant. For *T. nigroviridis*, the short reads were assembled by CLC genomics workbench using “Auto” parameters, then short reads were mapped to the assembled contigs. TPM was used for normalized expression values. The TPM values of *T. nigroviridis* were normalized with that of the grass puffer using TPM values of *GAPDH* gene. The average TPM values of *Hsp90, Hsp47, Hsp9*, and *unc45b* were compared using a t-test with Bonferroni correction.

### Phylogenetic, principal component, and ADMIXTURE analyses

The mapping data of eight grass puffers as well as two outgroup *Takifugu* species (three individuals each) were exported in bam file format and sorted and indexed using samtools^25^. The duplicated reads in bam files were marked by the MarkDuplicates algorithm in GATK v4.2 (https://gatk.broadinstitute.org/hc/en-us). We performed genotype calling using the HaplotypeCaller algorithm in GATK v4.2. Genotypes of all individuals were output as gvcf format (-ERC GVCF option). All gvcf files were combined into a single gvcf format file by the CombineGVCFs algorithm in GATK v4.2 and filtered by the VariantFiltration algorithm in GATK v4.2 with default parameters. The combined file was genotyped by the GenotypeGVCFs algorithm in GATK v4.2. We removed all indels, singleton, and doubleton sites to eliminate PCR and sequencing errors, and extracted bi-allelic sites with coverage equal to or more than five in all individuals and with GQ values equal to or more than twenty in all individuals.

The variants in a vcf file were converted to PHYLIP format. 10 kb sequences from the 5’ end of the PHYLIP format file were extracted and a model for Maximum Likelihood method was selected using MEGA ver. X ^26^. A phylogenetic tree was constructed using the Maximum Likelihood (ML) method using PhyML ver. 3.2 ^27^ with a model selection option “-m GTR” and with 100 bootstrap replications (Fig. 1: 116,941 sites).

We selected the Takifugu species from a gvcf file and filtered with the same condition described above, and performed a principal component analysis (PCA) using PLINK ver. 1.9 ^28^ with an option “--indep-pairwise 50 10 0.1” to explore the genetic relationship among all individuals (Figure 2A: 275155 sites).

ADMIXTURE ver. 1.3 ^29^ was run on the same dataset (Figure 2B: 275,155 sites) assuming 2 to 6 clusters (K=2-6).

## Acknowledgement

This work was supported by JST SPRING, Grant Number JPMJSP2104.

## Data availability statement

The nucleotide sequences were deposited in the DDBJ Sequenced Read Archive under a Bioproject PRJDB18283.

## Notes

### Competing Interest Statement

The authors have declared no competing interest.

### Summary of Updates

We added several new data. We also changed the format of our manuscript.

