## Supplemental data for "Adaptation of the eyes of grass puffer (*Takifugu niphobles*) to the riverine and marine environments"

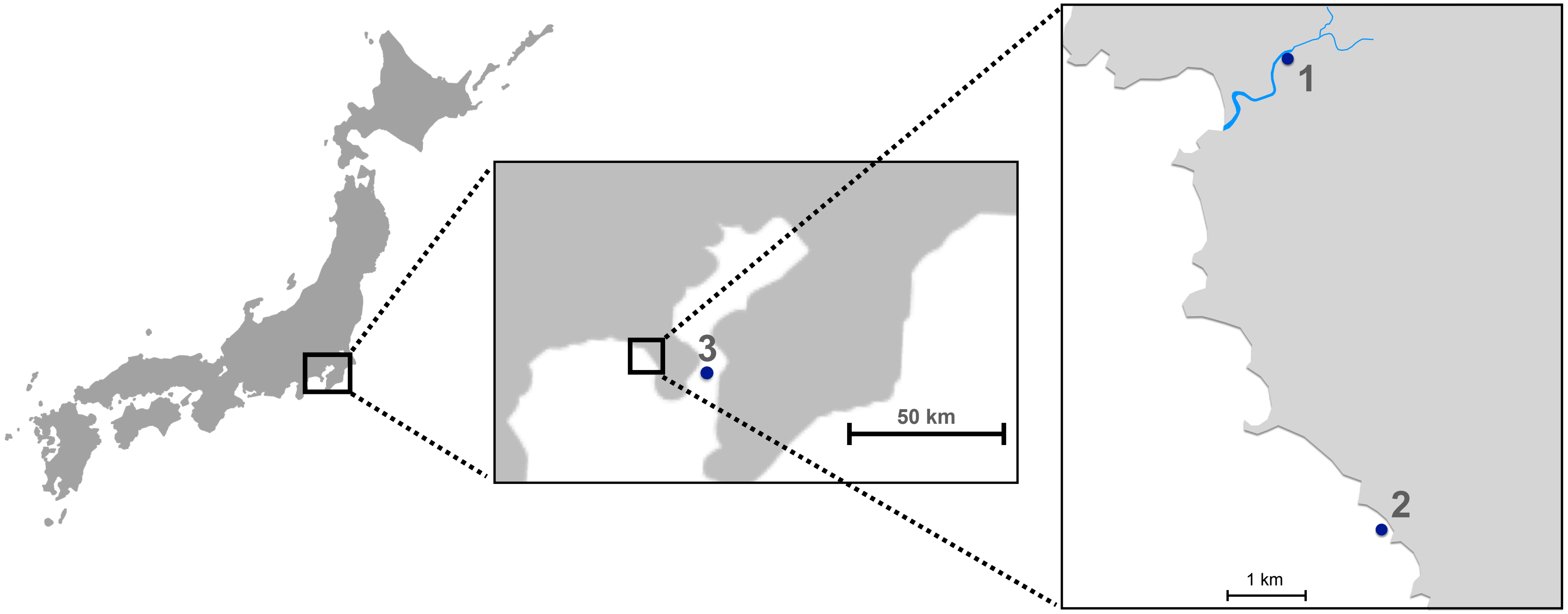


Fig. S1 Sampling locations
Fresh water and marine populations of *T. niphobles* were collected 1.3 km upstream from the mouth of the Tagoe River (1) and a beach (2) in Tateishi Park, Miura peninsula, respectively. Marine populations of *T. rubripes* and *T. synderi* were collected from Tokyo Bay (3).

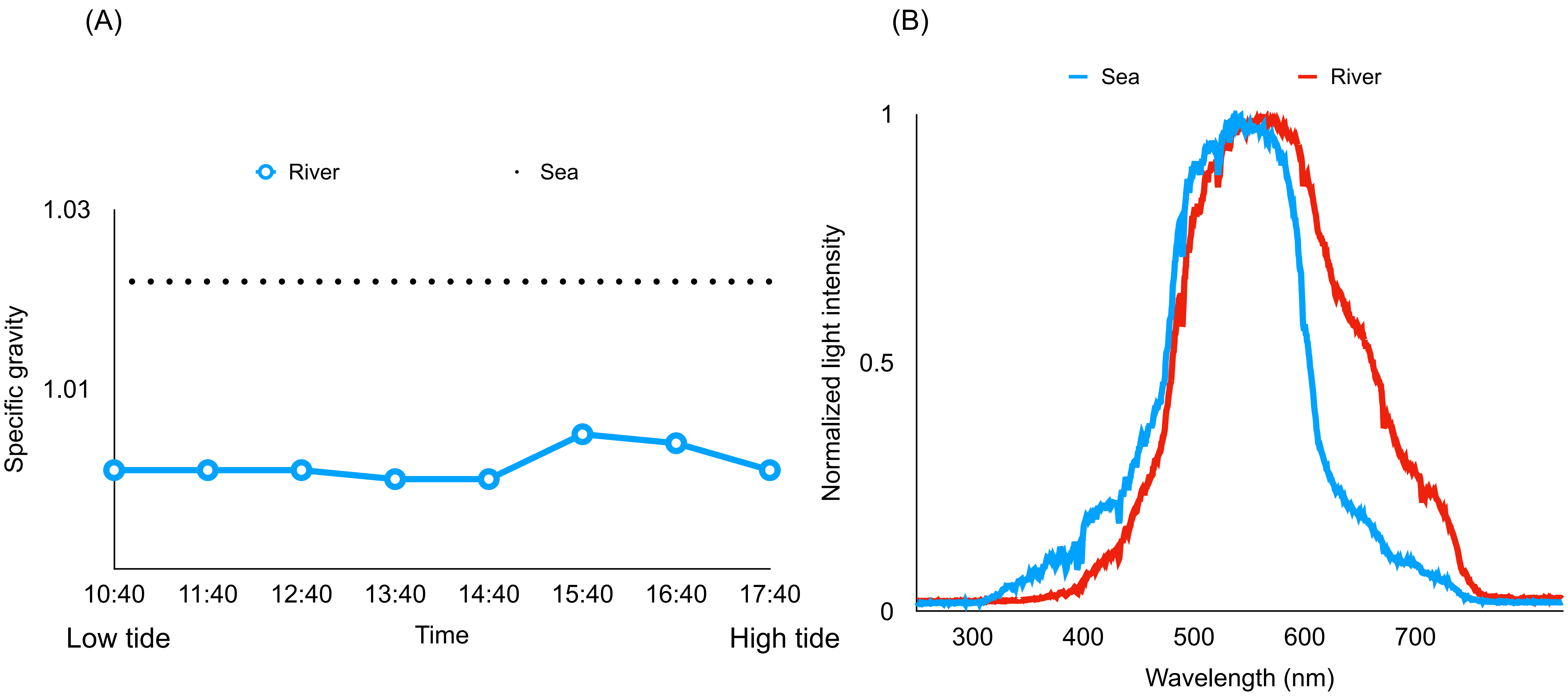


Fig.S2 The environments in the ocean and river at locations 1 and 2 in Fig. S1.
(A) The river's salinity was measured from low to high tide every hour. We measured the salinity in the sea only once.
(B) The light intensities in the river (red) and the sea (blue.

Table S1  Sample information

| Species | ID | Sampling location* | Environment | Sequence data | Mitochondrion Per identify (%) | Length (bp) |
|---|---|---|---|---|---|---|
| *Takifugu niphobles* | kr2 | 1 | River, depth 0-1 m* | DRX560745 | 99.32 | 16422 |
| | kr3 | 1 | River, depth 0-1 m* | DRX560746 | 99.51 | 16446 |
| | kr4 | 1 | River, depth 0-1 m* | DRX560747 | 99.56 | 16444 |
| | kr5 | 1 | River, depth 0-1 m* | DRX560748 | 99.55 | 16450 |
| | ks1 | 2 | Marine, depth 0-3 m* | DRX560749 | 99.57 | 16444 |
| | ks2 | 2 | Marine, depth 0-3 m* | DRX560750 | 99.43 | 16450 |
| | ks4 | 2 | Marine, depth 0-3 m* | DRX560751 | 99.51 | 16445 |
| | ks5 | 2 | Marine, depth 0-3 m* | DRX560752 | 99.55 | 16444 |
| *Takifugu rubripes* | tora5 | 3 | Marine, depth 5-15 m* | DRX560753 | 99.82 | 16447 |
| | tora6 | 3 | Marine, depth 5-15 m* | DRX560754 | 99.42 | 16449 |
| | tora7 | 3 | Marine, depth 5-15 m* | DRX560755 | 99.56 | 16424 |
| *Takifugu poecilonotus* | kom1 | 3 | Marine, depth 5-15 m* | DRX560756 | 99.79 | 16448 |
| | kom2 | 3 | Marine, depth 5-15 m* | DRX560757 | 99.69 | 16353 |
| | kom3 | 3 | Marine, depth 5-15 m* | DRX560758 | 99.76 | 16445 |
| *Canthigaster rivulata* | kitaF | 2 | Marine, depth 0-3 m* | DRX560763 | 96.59 | 16448 |
| *Tetraodon nigroviridis* | tn1 | - | River** | DRX560759 | 98.83 | 7090 |
| | tn2 | - | River** | DRX560760 | 98.84 | 7065 |
| | tn3 | - | River** | DRX560761 | 98.83 | 7078 |
| | tn4 | - | River** | DRX560762 | 99.22 | 7069 |

*Locations are shown in Fig. S1

**Based on Rainboth 1996

Table S2 Differentially expressed genes between riverine and marine populations

| Gene name | Fold change | FDR p-value |
|---|---|---|
| **heat shock protein 9** | **11.63512** | **9.14E-12** |
| **heat shock protein 47** | **8.033372** | **0.000104** |
| **unc45b** | **7.199732** | **0.000111** |
| dbpb | -6.41785 | 0.001459 |
| **TMX-3** | **39.99056** | **0.001459** |
| HMG-T1 | -5.44302 | 0.00178 |
| fosb | -5.99245 | 0.003654 |
| PCDH-1 | -18.4228 | 0.004062 |
| PAR bZIP | -4.76829 | 0.004062 |
| uncharacterized protein | -7.22996 | 0.007466 |
| uncharacterized protein | -16.0397 | 0.012557 |
| galectin-3-binding protein A | -29.925 | 0.019941 |
| f13a1 | -58.2615 | 0.019941 |
| uncharacterized protein | -9.99593 | 0.022636 |
| **heat shock protein 90** | **5.165485** | **0.024001** |
| CYP24A1 | -7.76695 | 0.03415 |
| CD82 | -7.80612 | 0.03415 |
| CABP5 | -4.35437 | 0.036797 |
| NLRC3 | -6.44915 | 0.040431 |
| SPC24 | -7.40586 | 0.040431 |
| uncharacterized protein | -5.51861 | 0.040431 |
| **FGA** | **6.7921** | **0.046341** |
| ADP-ribosylation factor | -7.30293 | 0.046341 |
| otoancorin | -10.91 | 0.046341 |
| immunoglobulin light chain | -11.4349 | 0.046341 |

Genes with higher expression levels in riverine grass puffer population are shown in bold.
